# The histone deacetylase activity of HDAC1/2 is required to safeguard zygotic genome activation in mice and cattle

**DOI:** 10.1101/2021.09.11.459880

**Authors:** Yanna Dang, Shuang Li, Panpan Zhao, Lieying Xiao, Lefeng Wang, Yan Shi, Lei Luo, Shaohua Wang, Huanan Wang, Kun Zhang

## Abstract

The genome is transcriptionally inert at fertilization and must be activated through a remarkable developmental process called zygotic genome activation (ZGA). The gene expression pattern formed over the course of ZGA is required for establishing totipotency in early embryos and subsequent development. Substantial epigenetic reprogramming contributes significantly to the pronounced change in gene expression during ZGA, however the mechanism has yet to be resolved. Here, we find histone deacetylase 1 and 2 (HDAC1/2) are critical histone modifiers that regulate ZGA through the histone deacetylase activity. Notably, we show that H3K27ac level declines dramatically during ZGA with a dynamic change in its genome-wide distribution. In mouse embryos, ectopic expression of HDAC1/2 dominant negative mutant leads to a failure of H3K27ac removal and a developmental arrest at 2-cell stage. RNA-seq results reveal a remarkable transcriptomic change with 6565 differentially expressed genes identified. Further analysis shows 64% of down-regulated genes are ZGA genes and 49% of up-regulated genes are developmental genes. Low input ChIP-seq analysis exhibits an increase and decrease of H3K27ac enrichment at the promoter region of up- and down-regulated genes, respectively. Moreover, HDAC1 mutants prohibited removal of broad H3K4me3 domain via impeding the expression of *Kdm5s* during ZGA. Importantly, the developmental block can be greatly overcome through injection of *Kdm5b* mRNA and expression of the majority of dysregulated genes partially corrected. Similar functional significance of HDAC1/2 in ZGA is conserved in bovine embryos. Together, we propose that HDAC1/2 is indispensable for mouse and bovine ZGA via creating correct transcriptional repressive and active states.

## INTRODUCTION

The highly differentiated germ cells (sperm and egg) go through the fertilization process to form a totipotent zygote, which marks the beginning of a new life. However, the new life must complete a developmental process called maternal zygotic transition (MZT) or oocyte-to-embryo transition (OET) to acquire its developmental independence^1, 2^. During this transition, the genome is transcriptional quiescent initially and the development control relies on maternal derived transcripts and proteins. These maternal factors gradually undergo clearance while zygotic genome is activated (ZGA). ZGA typically occurs in two consecutive waves, termed minor and major ZGA. The timing of ZGA varies among mammals. In mice, minor and major ZGA occurs in late one-cell and mid-to-late 2-cell embryos, respectively. The gene expression pattern established during ZGA is required for setting up totipotency in early embryos and the ensuing development. Therefore, a central question in biology is the defining feature of chromatin states during ZGA and how it contributes to changes in gene expression.

A considerable epigenetic reprogramming is a remarkable feature of early stage embryogenesis^3, 4, 5^. In particular, DNA methylations and various histone modifications are subject to dramatic remodeling, which is believed to ensure correct gene expression program and ZGA. Indeed, aberrant DNA methylation and histone modifications in fertilized or cloned embryos have been closely linked to abnormal gene expression and developmental failure^6, 7^. However, studies of molecular mechanisms of epigenetic reprogramming is traditionally limited by the scarce samples and a lack of chromatic analysis tools in early embryos.

With the rapid advance of low-input chromatin analysis methods including micro-scale ChIP-seq and CUT&RUN^8, 9, 10, 11, 12, 13^, how histone modifications patterns are passed, reprogrammed, and established from oogenesis to early embryogenesis have been unveiled and their correlation with gene expression profiles determined^8, 9, 10, 14^. However, the causal relationship between these histone modifications and gene expression as well as the crosstalk between histone modifications have yet to be resolved.

Histone acetylation is widely considered as a hallmark of active gene expression^15^. As one well-characterized histone acetylation, H3K27ac is associated with increased release of RNA polymerase II into active transcription and is typically accumulated at promoters and enhancers of active expressed genes^16^. In zebrafish embryos, there is a global loss of H3K27ac marks at enhancers while widespread de novo H3K27ac is acquired at promoters prior to ZGA^17^. Moreover, loss of H3K27ac leads to abnormal ZGA and results in early embryonic mortality^17^. In *Drosophila* embryos, ectopic H3K27ac prematurely activate lineage-specific genes at ZGA and causes embryonic lethality^18^. Therefore, these results indicate the importance of H3K27ac reprogramming in early embryonic development. However, the dynamics of H3K27ac and its regulatory mechanism in mammalian embryos remains largely unclear. This question is particularly important since growing evidence indicate chromatin reprogramming is quite different among mammals^19, 20^.

Histone acetylation is directly controlled by histone acetyltransferases (HATs) and histone deacetylases (HDACs)^21^. Mutations of these enzymes often lead to embryonic lethality in mice and are connected with human diseases^21^. HDAC1 is the first HDAC to be characterized. To date, four classes of 18 HDACs have been identified in mammals. Among them, class I family members (HDACs 1, 2, 3 and 8) display highest sequence homologous to HDAC1 and ubiquitously expressed, localized in the nucleus, and exhibit high catalytic activity towards histone substrates. HDAC1 and HDAC2 are the top expressed HDACs in mouse oocytes and embryos^15^. Genetic evidence reveals an overlapping and essential role of HDAC1 and 2 during oogenesis^22^. Likewise, our recent work established a redundant role of HDAC1 and HDAC2 in mouse early embryogenesis by employing siRNA-mediated gene silencing approach^23^. However, the specific role of HDAC1/2 in ZGA remains unclear given the technical difficulty in acutely removing maternal HDAC1/2.

In this study, we demonstrate that HDAC1/2-mediated removal of H3K27ac is required for ZGA in both mice and cattle by using a combination of dominant negative approach and pharmacological approach. HDAC1/2 mutants lead to a large-scale dysregulation of gene expression during ZGA and a developmental arrest at 2-cell stage. Mechanistically, HDAC1 mutants results in a deviant genome-wide distribution of H3K27ac and prohibits the removal of broad H3K4me3 domain by disrupting the expression of *Kdm5s*. Moreover, the developmental phenotype can be greatly rescued through injection of *Kdm5b* mRNA. We propose that HDAC1/2 are critical histone modifiers that maintain the transcriptional states during ZGA through their histone deacetylase activity.

## RESULTS

### A global loss of H3K27ac prior to major ZGA in mice and cattle

To probe the specific role of H3K27ac in ZGA, we first measured the level of H3K27ac in mouse and bovine early embryos around the time when ZGA occurs by immunofluorescence (IF). In mice, relatively strong signals of H3K27ac were detected in both maternal and paternal pronucleus at zygote stage and its nuclear intensity drastically diminished from the early and late 2-cell stages, concurrent with mouse major ZGA (Fig 1a). In cattle, H3K27ac is still relatively bright at 4-cell stage, when minor ZGA occurs, however, it underwent a sharp decrease at 8- and 16-cell stage, coinciding with bovine major ZGA (Fig 1b). These data are in agreement with previous studies in mice^24^ and such coincidence with ZGA was also reported in pigs^25^, suggesting that this global loss of H3K27ac is conserved in mammals.

**Fig. 1.**
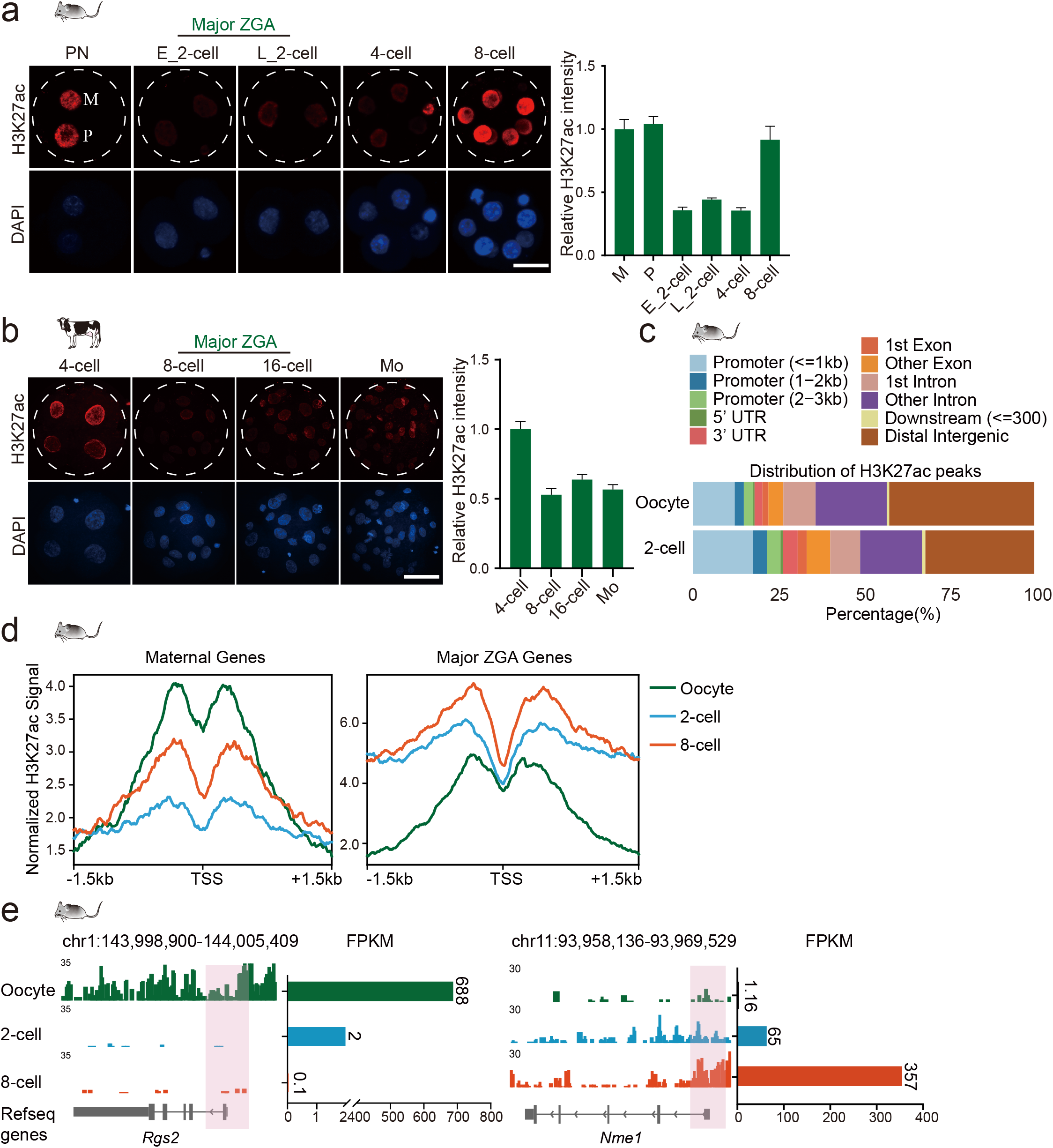
A global loss of H3K27ac prior to major ZGA in mice and cattle. **a,** Immunofluorescence staining of H3K27ac in mouse pronuclear stage (PN) zygote, early 2-cell (E_2-cell), late 2-cell (L_2-cell), 4-cell and 8-cell embryos. Left: representative images; M, maternal pronucleus, P, paternal pronucleus; scale bar, 25 μm. right: H3K27ac intensity relative to maternal pronucleus. Three replicates were conducted with at least 10 embryos in each stage, and data shown as Means ± SEM. **b,** Immunofluorescence staining of H3K27ac in bovine 4-, 8-, 16-cell embryos and morulae. Left: representative images; scale bar, 25 μm. right: H3K27ac intensity relative to 4-cell embryos. **c,** Feature distribution of H3K27ac peaks in oocyte and 2-cell embryos. **d,** Profiles of H3K27ac signal at promoters of maternal (left) and major ZGA (right) genes in mouse oocytes and preimplantation embryos. The H3K27ac signal is normalized to Reads Per Kilobase per Million mapped reads (RPKM) by DeepTools. **e,** Browser snapshot of H3K27ac distribution and Fragments Per Kilobase per Million mapped reads (FPKM) of RNA-seq at a representative maternal gene (left) and ZGA gene (right).

The immunostaining results were only quantitative for the global H3K27ac levels and didn’t reflect the changes of its exact chromatin distribution. To look into the dynamics of genome-wide distribution of H3K27ac around ZGA, we compared the H3K27ac map from oocyte to 2-cell stage using the published low-input ChIP-seq data^8^ and observed a dramatic change of H3K27ac across the genome. In particular, the global distribution of H3K27ac peaks at promoter regions was accumulated while the one at distal regions was decreased (Fig 1c). Because the chromatin feature of enhancer is not established during oogenesis and early embryogenesis, H3K27ac’s enrichment at enhancer regions is not analyzed in the present study. However, the dynamic changes in distal H3K27ac suggest a reduction of H3K27ac enrichment in enhancers. In sum, the coincidence of H3K27ac loss and major ZGA suggests a functional connection of H3K27ac reprogramming with ZGA in mammals.

To explore the relationship between H3K27ac and gene expression during ZGA, we first classified genes by their relative expression pattern from oocyte to late 2-cell stage embryos into three clusters: maternal, minor ZGA, and major ZGA by using the published RNA-seq data^26^ (Fig S1a). Then we generated H3K27ac landscape for these genes^8^, and observed an accumulation of H3K27ac at the active promoters (major ZGA genes) from oocyte to 2-cell stage embryo while the level of H3K27ac in those inactive promoters (maternal genes) decreased (Fig 1d). Moreover, individual browser snapshot of H3K27ac displayed a positive correlation between H3K27ac enrichment and gene expression in oocytes and early embryos (Fig 1e). As comparison, we calculated the distribution of H3K27me3 and H3K4me3 at promoters of these genes^9, 10^ (Fig S1b and S1c) and found no correlation between their enrichment and gene activation. Thus, H3K27ac is a reliable marker to distinguish active and repressed promoters in mouse 2-cell stage embryos. Overall, these results suggest the large-scale reprogramming of H3K27ac plays a critical role in safeguarding the correct gene expression profile during ZGA in mammals.

### The histone deacetylase activity of HDAC1/2 is required for H3K27ac loss and development during ZGA

To interrogate if the decrease of H3K27ac at ZGA depends on embryonic transcription, we blocked transcriptional activity by treating mouse zygotes with α-amanitin. As anticipated, embryos with α-amanitin treatment were arrested at 2-cell stage. IF analysis revealed that H3K27ac level is greater in α-amanitin-treated embryos (Fig 2a), suggesting the H3K27ac decrease is transcription-dependent.

**Fig. 2.**
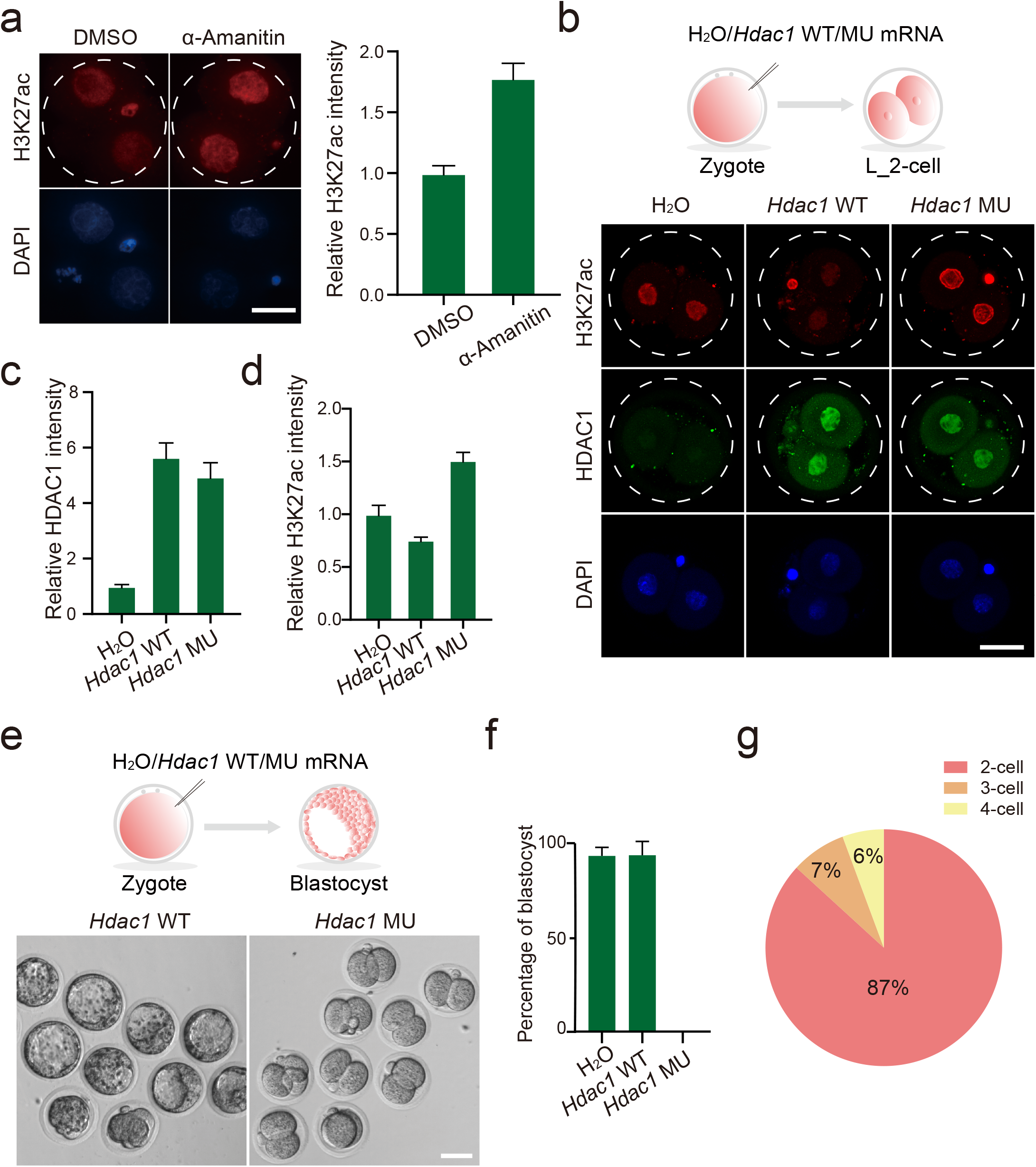
The histone deacetylase activity of HDAC1/2 is required for H3K27ac loss and development during ZGA. **a,** Immunofluorescence staining of H3K27ac in mouse late 2-cell embryos treated with α-Amanitin (10 μg mL^-1^) and DMSO (negative control). Left: representative images, scale bar, 25 μm, right: H3K27ac intensity relative to DMSO-treated embryos. **b,** Experimental scheme (top) and immunofluorescence staining (bottom) of HDAC1 and H3K27ac in late 2-cell embryos. **c,** HDAC1 intensity relative to embryos injected with H_2_O. **d,** H3K27ac intensity relative to embryos injected with H_2_O. **e,** Representative images of embryos injected with *Hdac1* WT mRNA and *Hdac1* MT mRNA at day 4.5 after fertilization. **f,** Percentage of blastocysts at day 4.5 after fertilization in three groups. **g,** Distribution of cleavage embryos found in embryos injected with *Hdac1* mutant mRNA at day 4.5 after fertilization.

We hypothesized that histone deacetylase is actively responsible for the removal of H3K27ac. Transcriptomic analysis revealed that Hdac1/*HDAC1* and Hdac2/*HDAC2* were the most abundant among all *Hdacs/HDACs* in both mouse (Fig S2a) and bovine early embryos (Fig S2b). In particular, *Hdac1* mRNA abundance increased gradually and visibly during ZGA in both species (Fig S2a and S2b). In accordance with the mRNA change, the nuclear intensity of HDAC1 was increased during ZGA (Fig S2c and S2d). These results suggest that the decline of H3K27ac is attributed to the increase of HDAC1/2.

To delve into the function of HDAC1/2 in ZGA, we treated the zygotes with FK228, a HDAC1/2-specific inhibitor^27^. Results confirmed a robust increase of H3K27ac level in FK228-treated embryos (Fig S2e and S2g). Embryo culture results showed FK228-treated embryos arrested at 2-cell stage in mice (Fig S2f). Concomitantly, FK228 treatment resulted in a developmental arrest in cattle and further cell number analysis with DAPI staining clarified that these embryos arrested at 8/16-cell stage (bovine ZGA; Fig S2h and S2i).

To probe the specific effects of the histone deacetylase activity of HDAC1, we generate a mutant in the histone deacetylase domain. The histone deacetylase domain conserved to all Class I HDACs consists of a stretch of more than 300 amino acids and a mutation in the amino acid sequence destroys its enzymatic activity^28^. We therefore constructed a HDAC1 mutant in which the residue histidine 141 (H141) was replaced by alanine^28^. The resultant mutant is referred to here as H1MU while the wild type is referred to as H1WT. We microinjected early zygotes with mRNA for H1WT and H1MU. Water was injected in control groups (Fig 2b). The mutation does not affect the nuclear localization of HDAC1 in early embryos. As expected, confocal microscopy analysis of H1WT and H1MU embryos showed an apparent increase in immunostaining of HDAC1 (Fig 2b and 2c). In comparison with control groups, there is a slight decrease of H3K27ac in H1WT embryos, further confirming the effectiveness of *Hdac1* mRNA injected. In contrast, H1MU led to a significant increase in the global H3K27ac level (Fig 2b and 2d), suggesting the mutation efficiently abolished the histone deacetylase activity in mouse embryos. Moreover, nearly all of the embryos in H1MU group suffered developmental arrest at 2-cell stage (87%) while the control embryos exhibit robust development with 80% developing to blastocyst stage (Fig 2e, 2f and 2g).

HDAC1 and 2 are two homologous histone deacetylases and exhibit functional redundancy in oogenesis and preimplantation development^22, 23^. We therefore asked if these two proteins play a redundant role during ZGA. *Hdac2* mutant was also generated by employing a mutation in the catalytic domain^29^. Likewise, the mutation also abolished the histone deacetylase activity of HDAC2 and caused embryonic arrest at 2-cell stage (Fig S2j and S2k). Overall, these data revealed that the acute removal of H3K27ac mediated by HDAC1/2 is crucial for ZGA in both mouse and bovine embryos.

### HDAC1 mutation results in aberrant gene expression pattern during ZGA

To examine if the histone deacetylase activity of HDAC1 affect the genome-wide transcriptional program, we then measured the level of phospho-Ser2 (Ser2P), a hallmark of RNA polymerase II activity, and found no visible difference between H1MU and H1WT embryos. This result suggests a locus-specific role of HDAC1 in transcriptional regulation.

To further identify the molecular targets of HDAC1 and gain insight into the mechanisms of developmental failure of H1MU embryos, we collected early and late 2-cell stage embryos from H_2_O, H1WT, and H1MU groups and performed RNA-seq (Fig. 3a). Two independent replicates for RNA-seq samples from each group displayed high correlation (Fig. S4a). In addition, *Hdac1* was significantly increased in both H1WT and H1MU groups, further confirming a robust overexpression efficiency (Table S2).

**Fig. 3.**
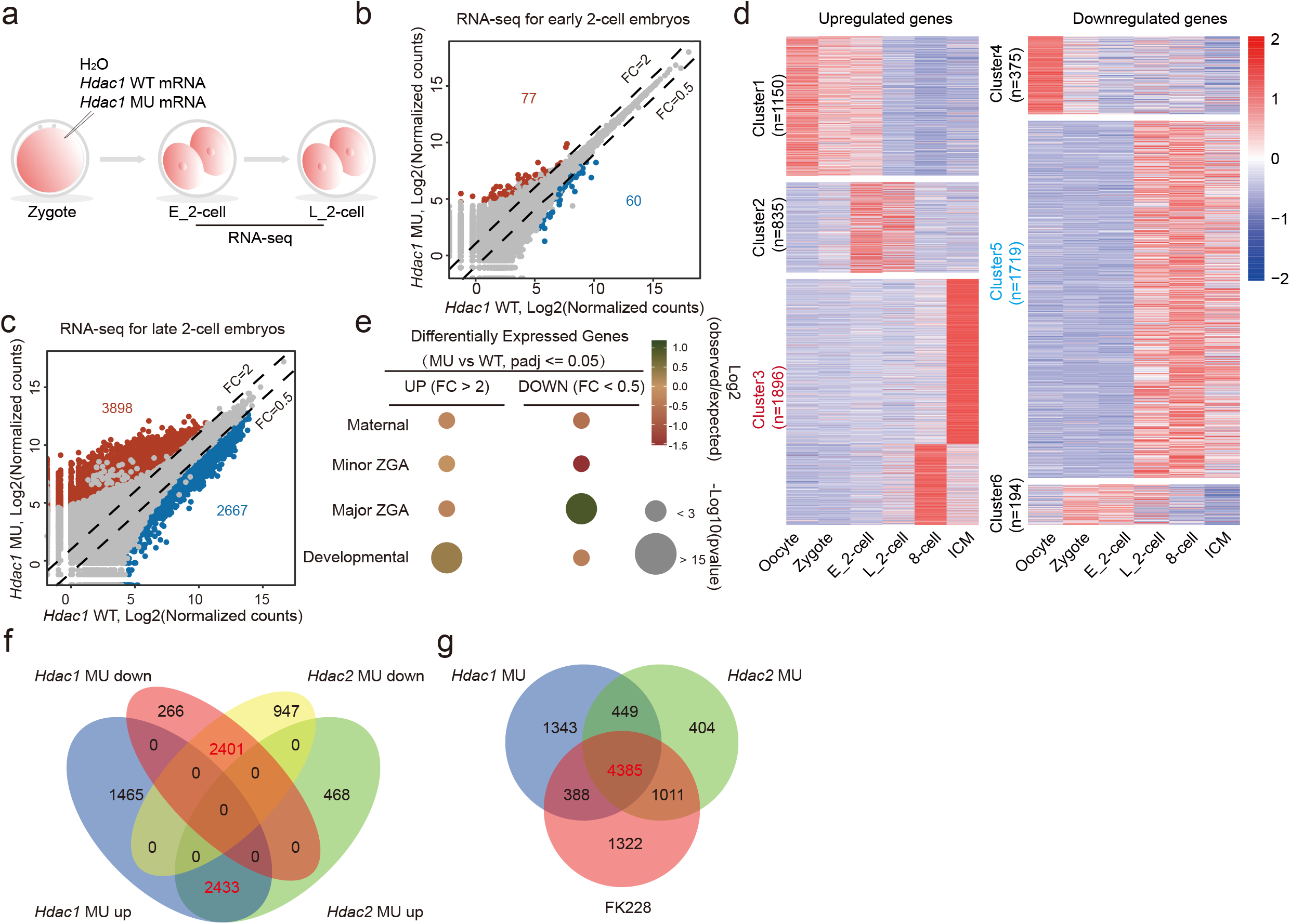
HDAC1 mutation results in aberrant gene expression pattern during ZGA. **a,** Experimental scheme of RNA-seq. **b and c,** Scatter plots showing global gene expression in embryos injected with *Hdac1* WT mRNA and *Hdac1* MU mRNA at early 2-cell stage (b) and late 2-cell stage (c). Two RNA-seq replicates are generated for differential expression analysis, and the read counts are normalized by DESeq2. Dash lines indicate the threshold of fold change (MU/WT), and grey dots refer to genes with *padj* > 0.05, while dots in red and blue refer to genes with *padj* < =0.05. Numbers of up- and down-regulated genes are indicated in the figures. **d,** Heatmaps showing relative expression of up- (left, *Hdac1* MU/WT FC >2 and *padj* <=0.05) and down-regulated (right, *Hdac1* MU/WT FC <0.5 and *padj* <=0.05) genes during mouse preimplantation development. The differential expressed genes are classified to 6 clusters by *k-means* clustering. **e,** Overlap of all differentially expressed genes with different gene categories. The gene categories are generated with *k-means* clustering of RNA-seq data for mouse oocytes and zygote, early 2-cell, late 2-cell, 8-cell embryos and ICM. The color of bubbles refers to log_2_ ratio of number of observed genes in the gene set relative to randomly expected frequencies, and the size of bubbles shows -log_10_ of Fisher-exact test *p* value. **f,** Venn diagram showing overlap of up- and down-regulated genes induced by overexpression of *Hdac1* and *Hdac2* MU mRNA. Note that size of the circles doesn’t reflect the number of genes. **g,** Overlap of all differentially expressed genes induced by overexpression of *Hdac1* or *Hdac2* MU mRNA and treatment of FK228. Size of the circles doesn’t equal to the number of genes.

H1MU has no significant impact on minor ZGA as the analysis revealed only 77 and 60 up-regulated and down-regulated genes, respectively (Padj<=0.05; Fold change>=2 or <= 0.5) at early 2-cell stage in H1MU versus H1WT groups (Fig 3b). Moreover, at late 2-cell stage, there are only 88 genes differentially expressed in WT groups compared with control groups, suggesting H1WT overexpression has minor effect on the transcriptome during ZGA (Fig S4b). In contrast, 6565 genes were differentially expressed in late 2-cell embryos of H1MU compared to H1WT groups with 3898 and 2667 up-regulated and down-regulated genes, respectively (Fig 3c and Table S2). These transcriptomic comparisons at the two different embryonic stages clearly indicated HDAC1 specifically regulate gene transcription when major ZGA occurs.

Gene Ontology (GO) enrichment analysis of up-regulated genes using DAVID revealed overrepresentation of genes involved in transcription and sequence-specific DNA binding (Table S3). Meanwhile, similar analysis of down-regulated genes showed that a significant number of genes were involved in transcription and RNA splicing, ribosome biogenesis and nucleotide metabolism, which are known markers of ZGA^30^ (Table S4). We also noticed a drastic reduction of the expression of *Brg1/Smarca4^31^* and *Nfya^32^*, two genes known for their putative function in regulating mouse ZGA.

To assess the molecular feature of HDAC1’s target genes, we then examine the expression pattern of the up-regulated genes and down-regulated genes during normal preimplantation development. We categorized the up-regulated and down-regulated genes into three main classes each by *k-means* clustering. Surprisingly, a considerable number of up-regulated genes are maternal (Cluster 1) and developmental genes (Cluster 3), which are supposed to be silent, while the majority of down-regulated genes are ZGA genes (Cluster 5; Fig 3d and 3e). These results suggest HDAC1’s enzymatic activity is required not only for transcriptional silencing but transcriptional activation during ZGA.

To address directly if there is functional redundancy for HDAC1/2 in regulating gene transcription, we compared differentially expressed genes (DEGs) caused by H1MU and H2MU. Results indicated that the majority of up-regulated and down-regulated genes are common to both groups (Fig 3f and S4e), suggesting a compensatory role of HDAC1/2 in transcriptional regulation. We also observed FK228 treatment resulted in similar changes in the transcriptome (Fig 3g and S4e), indicating its specificity in inhibiting the histone deacetylase activity of HDAC1/2. In agreement with the result of H1MU, we observed the majority of up-regulated and down-regulated in H2MU or FK228-treated groups are maternal/developmental genes and ZGA genes, respectively (Fig S4f, S4g, and S4h).

### HDAC1 mutation results in aberrant distribution of H3K27ac in early embryos

To explore how H1MU affects the transcriptional program, we performed Ultra-Low-Input-NChIP-seq (ULI-NChIP-seq) of H3K27ac at late 2-cell stage and obtained high-quality reproducible data (Fig 4a, 4b, and S5a). The protocol used here is highly efficient since our data in control groups faithfully recapitulates the previously published H3K27ac ChIP-seq data^8^ with a high correlation (R=0.80; Fig S5b and S5c). Genome distribution of H3K27ac changed significantly in H1MU embryos. In particular, the H3K27ac enrichment at promoter and distal regions were decreased and increased, respectively (Fig 4c). By scanning the genome with a 5 kb sliding window, we identified H3K27ac-lost and -gained regions in H1MU embryos. In consistent with our IF results, H3K27ac-gained regions covered more genome than H3K27ac-lost regions (Fig 4d).

**Fig. 4.**
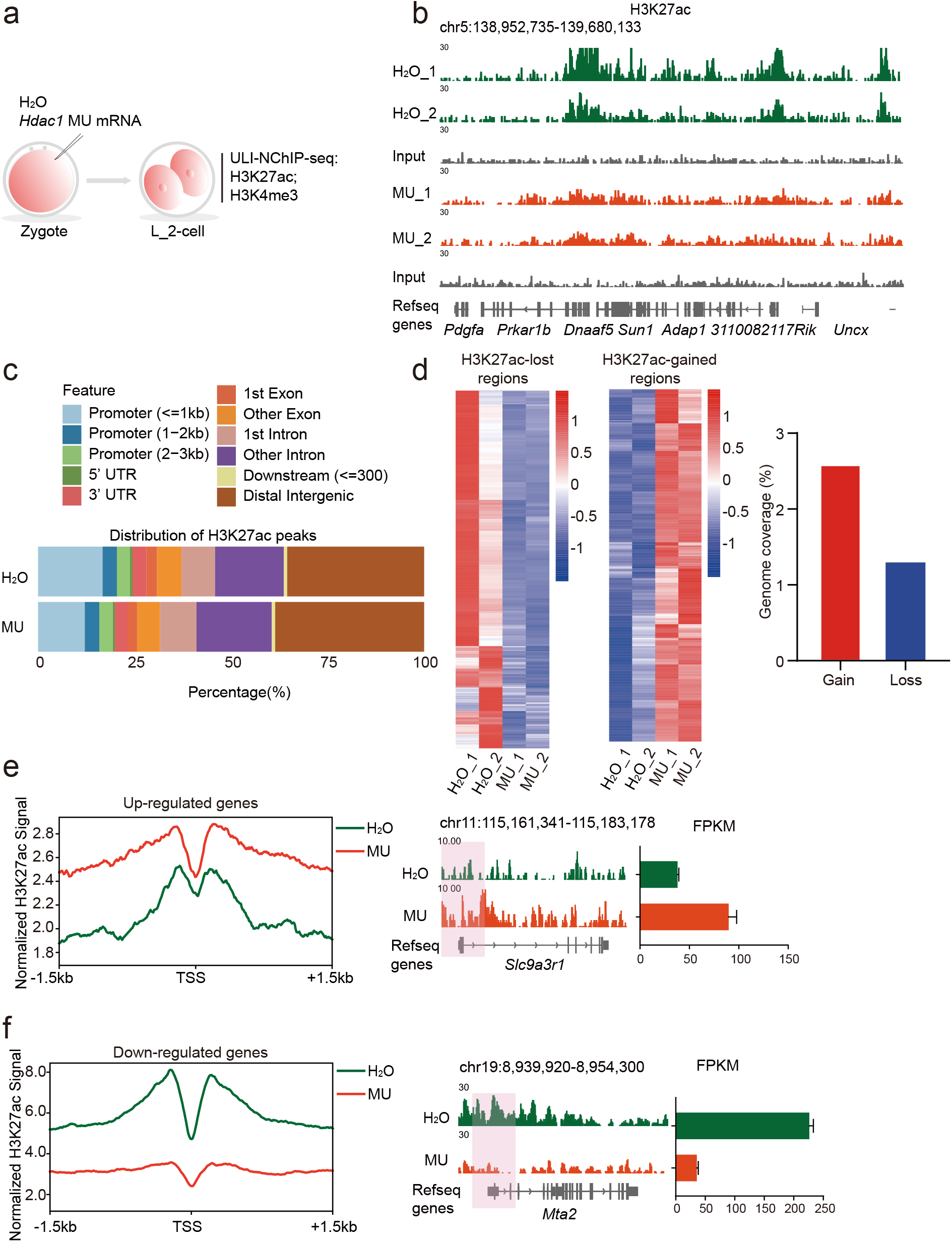
HDAC1 mutation results in aberrant distribution of H3K27ac in early embryos. **a,** Experimental scheme for ULI-ChIP-seq. **b,** IGV browser snapshots showing H3K27ac signals in two biological replicates of embryos injected with H_2_O and *Hdac1* MU mRNA respectively. **c,** Feature distribution of H3K27ac peaks in the two groups. **d,** Heatmaps showing z-score normalized RPKM of H3K27ac-lost and -gained regions. Bar plots display the genomic coverage for those regions. Identification of H3K27ac-lost and -gained regions are described in the Methods section. **e and f**, The left panel showing profile of H3K27ac signal at promoters of all up-regulated genes (*Hdac1* MU/WT FC>2 and padj <=0.05) (e) and down-regulated genes (*Hdac1* MU/WT FC <0.5 and padj <=0.05)(f). H3K27ac signal is normalized to RPKM by DeepTools. The right panel showing IGV browser snapshot of up-regulated gene (e) and down-regulated gene (f). The corresponding FPKM in RNA-seq data is showed in bar plots.

To establish if the epigenetic changes associated with the transcriptomic differences, we profiled H3K27ac signal in promoters of up-regulated and down-regulated genes. H3K27ac was accumulated in both promoters and gene bodies of up-regulated genes in H1MU group (Fig 4e and S5d). On the contrary, the promoters and gene bodies of down-regulated genes displayed reduced H3K27ac signal (Fig 4f and S5e). These results suggest that the disorder of gene expression pattern caused by H1MU is likely attributed to the aberrant H3K27ac distribution.

### DUX is enriched in H3K27ac-gained regions upon HDAC1 mutation

To identify sequence motifs for DNA-binding proteins that were potentially enriched in aberrant H3K27ac-deposited regions, we scanned the entire genome within a 1 kb window to calculate H3K27ac signals, and annotated motifs at the top 2% H3K27ac-increased and -decreased regions by HOMER. Notably, DUX was significantly enriched in the H3K27ac-increased regions (Fig 5a). However, these motifs were not detected in H3K27ac-decreased regions. Altogether, these results suggest that H1MU influences expression of active and repressed genes through distinct regulatory mechanisms.

**Fig. 5.**
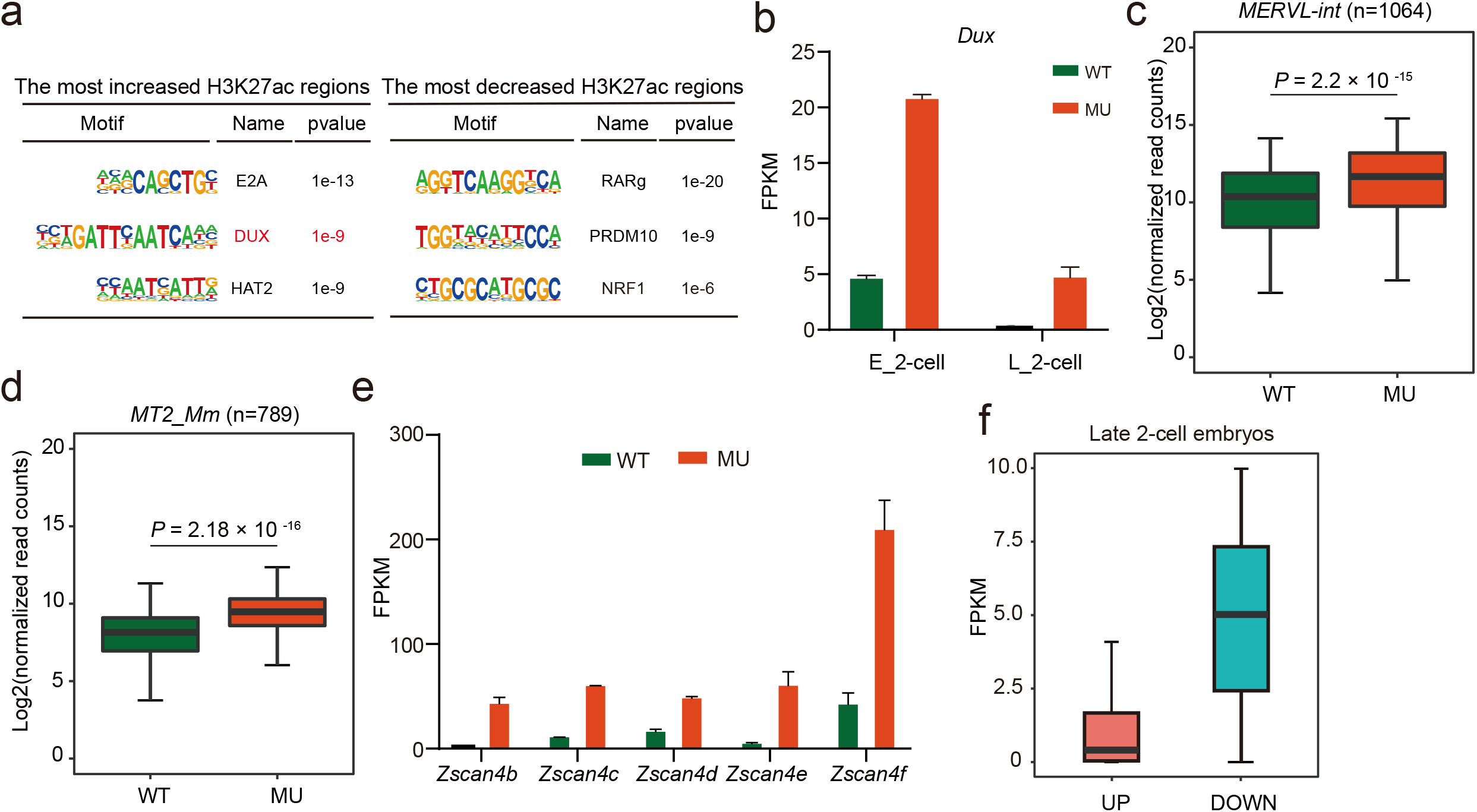
DUX is enriched in H3K27ac-increased regions upon HDAC1 mutation. **a,** Representative motifs enriched at the most H3K27ac-increased and -decreased regions (with *p* value) are listed. **b,** Relative expression of *Dux* (also known as *Duxf3*) in early and late 2-cell embryos. **c, d,** Box plots showing expression of *MERVL-int* (n=1064) and *MT2_Mm* (n=789) in late 2-cell embryos. Read counts are normalized by DESeq2. *P* values are calculated by Wilcoxon rank sum test. **e,** Relative expression of *Zscan4s* in late 2-cell embryos. **f,** Box plots showing average expression levels of up-regulated (*Hdac1* MU/WT FC>2 and padj <=0.05) and down-regulated (*Hdac1* MU/WT FC <0.5 and padj <=0.05) genes in H_2_O injection embryos at late 2-cell stage.

DUX (DUX4 in humans) is a putative transcriptional factor involved in regulation of mammalian ZGA^33, 34, 35^. *Dux* is supposed to be transiently expressed during early to mid-2-cell stage and rapidly disappears at late 2-cell stage (Fig 5b)^36^. However, although there was a distinct decrease of *Dux* in late 2-cell embryos of H1MU groups, the amount was still greater than that in control embryos (Fig 5b). *Zscan4s* and *ERVLs* are identified as target genes of DUX, and can be robustly expressed in mouse embryos overexpressing DUX^36^. We found the expression of *Zscan4s* and *ERVLs (MERVL-int and MT2_mm)* was also increased in H1MU embryos (Fig 5c, 5d and 5e). Therefore, it is highly possible that the insufficient repression of *Dux* is one leading factor causing the up-regulation of repressed genes, including developmental and maternal genes.

It has been reported that DUX4 function as a DNA-binding protein that recruit EP300/CBP through its C-terminals to activate expression of nearby genes^37^. Intriguingly, the average mRNA abundance of up-regulated genes in late 2-cell H1WT embryos is lower than those of down-regulated genes (Fig 5f), suggesting the timely loss of DUX in late 2-cell embryos is critical to maintain the low expression of those up-regulated genes.

### HDAC1 mutation inhibits the removal of broad H3K4me3 domain during mouse ZGA

H3K4me3 is an established hallmark of permissive promoters that can tether chromatin remodelers and transcriptional regulators. H3K27me3 is prevalently enriched in the promoters of repressed genes. We sought to investigate the potential functional relationship between H3K27ac and H3K4me3/H3K27me3. We first obtained genes marked by only H3K4me3 or H3K27me3 at 2-cell stage from the published ChIP-seq data^9^ and observed H3K27ac is accumulated at H3K4me3-marked promoters but not H3K27me3-marked promoters (Fig S7a). To further determine the colocalization of H3K4me3 and H3K27ac, we performed H3K4me3 ULI-NChIP-seq at late 2-cell stage and found more than 50% of H3K27ac peaks overlapped with those of H3K4me3 in control embryos (Fig 4a and 6a). The colocalization of H3K4me4 and H3K27ac prompts us to address the crosstalk between these two histone modifications.

**Fig. 6.**
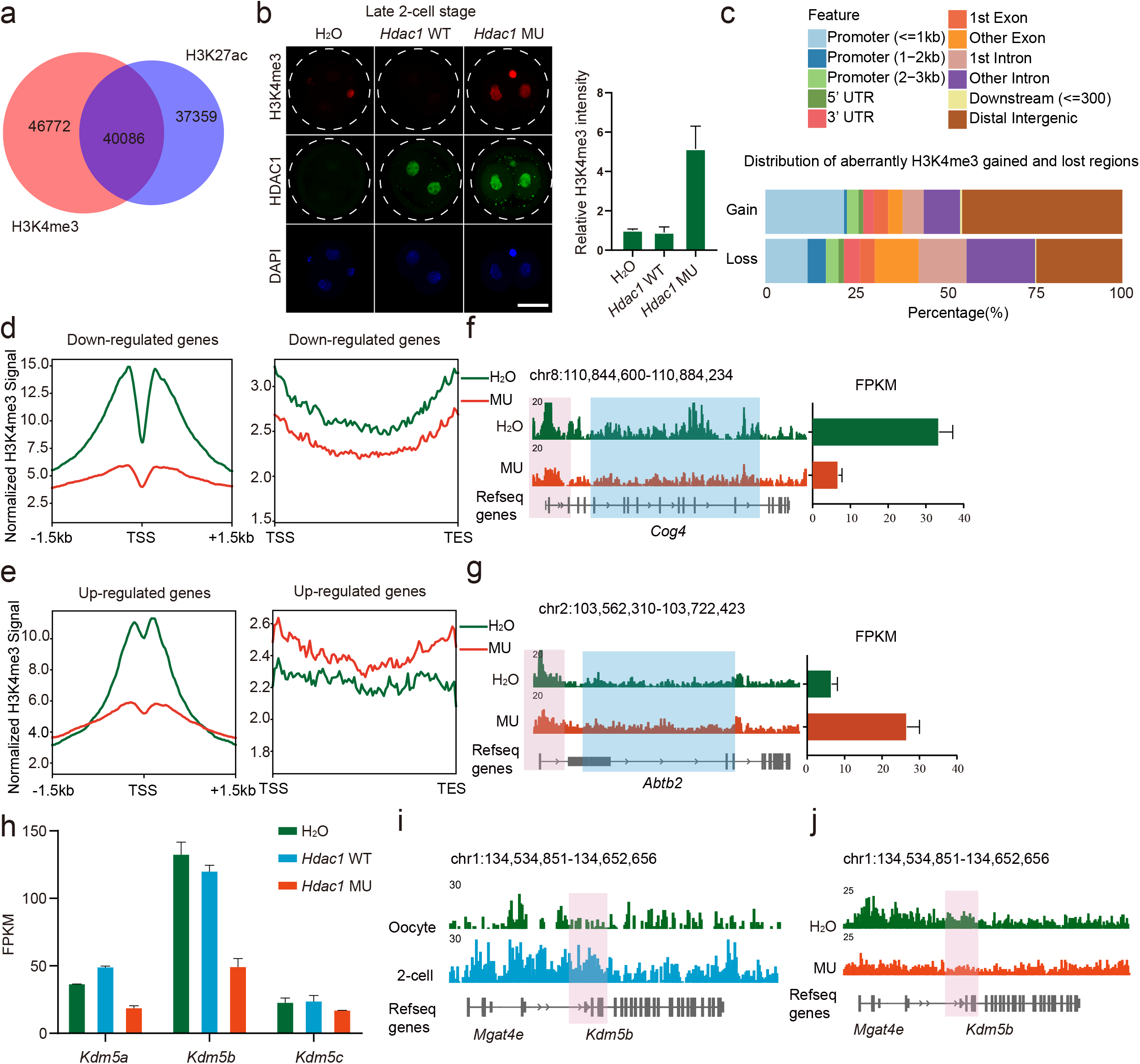
HDAC1 mutation inhibits the removal of H3K4me3 during mouse ZGA. **a,** Venn diagram showing the overlap of H3K27ac peaks and H3K4me3 peaks in mouse late 2-cell embryos of H_2_O injection group. **b,** Immunofluorescence staining of H3K4me3 and HDAC1 at late 2-cell stage in three groups. Left: representative images, scale bar, 25 μm, right: H3K4me3 intensity relative to H_2_O-injected embryos. **c,** Feature distribution of H3K4me3-gained and -lost regions. Identification of H3K4me3-lost and -gained regions are described in the Methods section. **d and e,** The left panel shows profiles of H3K27ac signal at promoters of all down-regulated genes (*Hdac1* MU/WT FC <0.5 and padj <=0.05) (d) and up-regulated genes (*Hdac1* MU/WT FC>2 and padj <=0.05) (e). The right panel shows profiles of H3K27ac signal in gene body regions of all down-regulated (d) and up-regulated (e) genes. H3K27ac signal is normalized to RPKM by DeepTools. **f and g,** Browser snapshots showing H3K27ac at down-regulated genes (f) and up-regulated genes (g). The relative expression is showed in bar plots. **h,** Relative expression of *Kdm5a, Kdm5b, Kdm5c* in late 2-cell embryos. **i,** Browser snapshots showing H3K27ac at *Kdm5b* in wildtype oocytes and 2-cell embryos. **j.** Browser snapshots showing H3K27ac at *Kdm5b* in late 2-cell embryos.

Genome-wide distribution of H3K4me3 is unique in early embryos for its broad domains established during oogenesis and removed upon ZGA^8, 9, 10^. We observed a notable level of H3K4me3 at early 2-cell stage in control groups and no difference was found in H1MU groups (Fig S6d). Furthermore, H3K4me3 signal can be barely seen in control embryos while the intensity was obviously increased in H1MU embryos, suggesting the removal of broad H3K4me3 domain is blocked (Fig 6b).

We then analyzed the global dynamics of H3K4me3 in H1MU and control embryos by ChIP-seq. We validated the H3K4me3 ChIP first and found there is high correlation between our data in control groups and the previously published H3K4me3 ChIP-seq data^10^ (R=0.81; Fig S6a and S6b). The two biological replicates for both groups exhibit high correlation, indicating a high reproducibility (Fig S6c and S6d). We sought to assess H3K4me3 signals at the most H3K27ac-increased and -decreased regions and found H3K4me3 was accumulated and removed correspondingly (Fig S7b and S7c). H3K4me3-gain regions accounted for nearly 50% at distal regions while H3K4me3-lost regions accounted for 25% at distal regions, further suggesting a block in removal of broad H3K4me3 domain (Fig 6c). In consistent with H1MU’s effect, we also found the erasure of broad H3K4me3 domain was suppressed when H2MU or FK228 treatment was employed (Fig S7e-g).

H3K4me3 reprograming is critical to ensure gene expression program during ZGA. We therefore sought to determine if H3K4me3 involved in transcriptomic changes in H1MU embryos. Surprisingly, H3K4me3 was reduced at promoters of both up- and down-regulated genes in H1MU group (Fig 6d-g). However, H3K4me3 signals were enhanced in gene body regions of up-regulated genes while it was reduced at gene body regions of down-regulated genes (Fig 6d-g), suggesting that the dysregulated gene expression could be ascribed to the sustained broad H3K4me3 domain. Altogether, HDAC1’s enzymatic activity is crucial for the immediate removal of broad H3K4me3 domain upon ZGA.

Broad H3K4me3 domains are removed by KDM5 family members during mouse ZGA^8, 9, 10^. Interestingly, GO analysis of down-regulated genes revealed genes involved in histone H3-K4 demethylation (Table S4). Indeed, mRNA abundance of all three *Kdm5s (Kdm5a, 5b, 5c)* was dramatically decreased in H1MU embryos (Fig 6h). *Kdm5b* is zygotic transcribed from early to late-2 cell stage (Fig S7g) while H3K27ac is accumulated at *Kdm5b*. In contrast, H1MU caused a reduction in H3K27ac enrichment at *Kdm5b*, suggesting H3K27ac regulates transcription of *Kdm5b*.

### KDM5B is a key mediator of the HDCA1 MU phenotype

We next asked if ectopic injection of *Kdm5* mRNA could rescue the H3K4me3 pattern and the developmental outcome of H1MU embryos (Fig 7a). Because *Kdm5b* normally begins its transcription at 2-cell stage, we thus performed microinjection of *Kdm5b* mRNA into two individual blastomeres of early 2-cell embryos. Results showed microinjection of *Kdm5b* resulted in a significant decrease of global H3K4me3 in H1MU embryos, indicating a recovery of H3K4me3 reprogramming (Fig 7b). As a consequence, the majority of the co-injected embryos could overcome the 2-cell block and develop beyond 4-cell stage (Fig 7c and 7d). RNA-seq analysis revealed that the expression of 77.17% of downregulated genes and 47.46% of upregulated genes in H1MU groups could be at least partially corrected by ectopic expression of *Kdm5b* (Fig 7e), and the rescued genes were significantly enriched in major ZGA (Fig 7f), suggesting that the accurate distribution of H3K4me3 along with H3K27ac is required for ZGA. Remarkably, expression of *Dux* and its targets *Zscan4s* dropped to normal levels when overexpressing *Kdm5b* (Fig 7g and 7h), suggesting that HDAC1 regulates DUX through KDM5B. Together, these findings implicate KDM5B as a key molecular mediator of HDAC1 function in controlling H3K4me3 reprogramming.

**Fig. 7.**
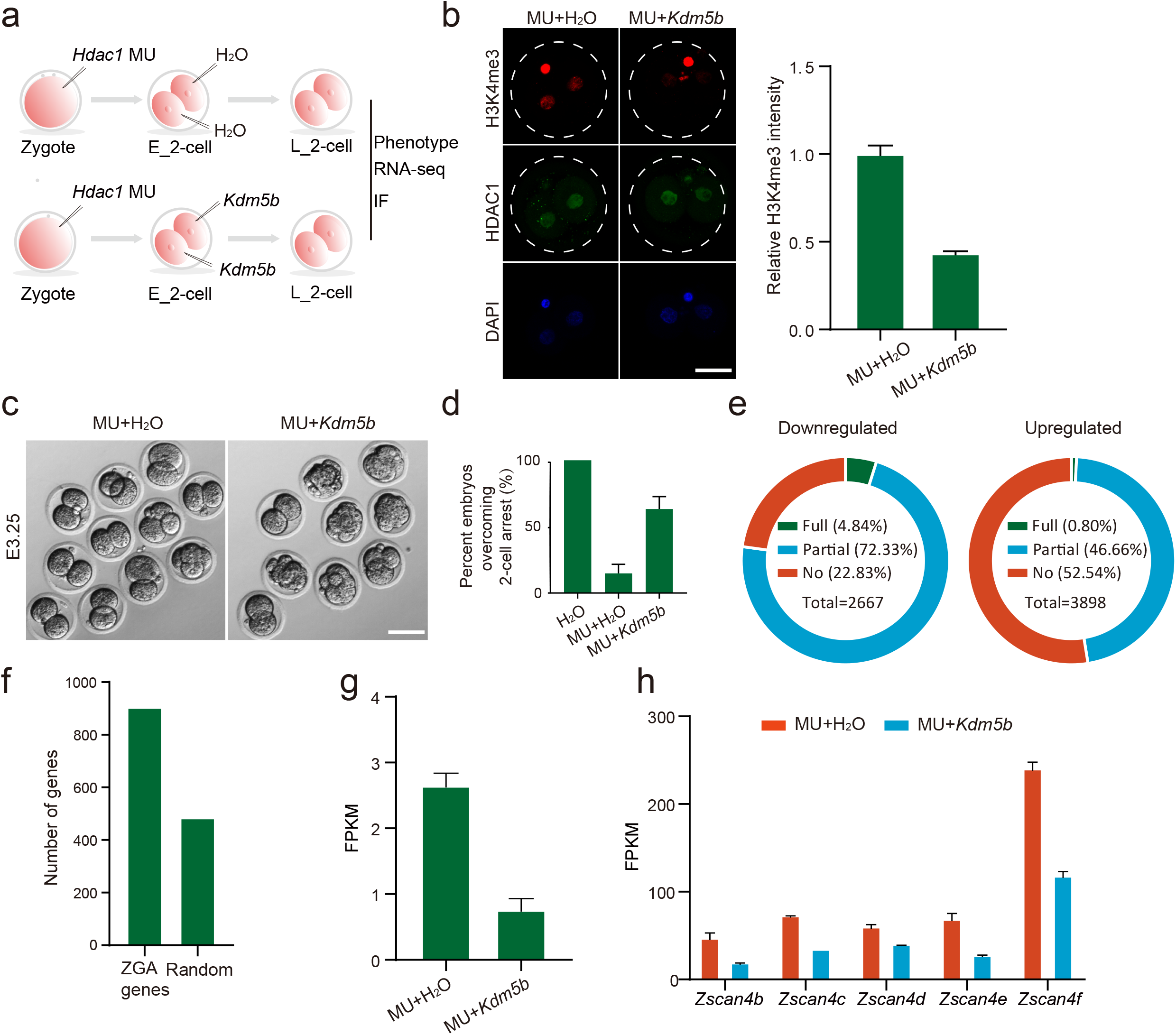
KDM5B is a key mediator of the HDCA1 MU phenotype. **a,** Experimental scheme showing overexpression of exogenous *Kdm5b* in early embryos. **b,** Immunofluorescence staining of H3K4me3 and HDAC1 at late 2-cell stage. Left: representative images, scale bar, 25 μm, right: H3K4me3 intensity relative to embryos injected with *Hdac1* MU mRNA and H_2_O. **c,** Developmental progression of embryos in the two groups at day 3.25 after fertilization. **d,** The ratio of embryos developed beyond 2-cell stage at day 3.25 after fertilization. **e,** Donut chart of down-regulated genes (*Hdac1* MU/WT, left) and up-regulated genes (*Hdac1* MU/WT, right) based on their extent of rescue in *Hdac1* MU mRNA and *Kdm5b* mRNA injection embryos. For down-regulated genes, full rescue means genes characterized by a log_2_ ratio of MU+*Kdm5b/Hdac1* WT ≥0; partial rescue refers to genes characterized by a log_2_ ratio of MU+*Kdm5b/Hdac1* MU >0; no rescue genes are characterized by a log_2_ ratio of MU+*Kdm5b/Hdac1* MU ≤0. For up-regulated genes, full rescue means genes characterized by a log_2_ ratio of MU+*Kdm5b/Hdac1* WT ≤0; partial rescue refers to genes characterized by a log_2_ ratio of MU+*Kdm5b/Hdac1* MU <0; no rescue genes are characterized by a log_2_ ratio of MU+*Kdm5b/Hdac1* MU ≥0. **f,** Number of up-regulated (MU+*Kdm5b*/ MU+H_2_O) major ZGA genes and the number expected by chance, Fisher exact test *P* value < 10^-10^. **g,** Relative expression of *Dux* (also known as *Duxf3*). **i,** Relative expression of *Zscan4s*.

### Function of HDAC1 on ZGA is conserved in mouse and bovine embryos

Recent studies in epigenome reprogramming have revealed remarkable differences among mammals^19, 38^. To determine if the effect of HDAC1 on transcriptional regulation during ZGA is conserved, we collected DMSO and FK228-treated 8-16 cell embryos and performed RNA-seq. Results showed 1990 and 2794 up-regulated and down-regulated genes, respectively (Fig 8a, 8b, S8a and S8b). 74% (2061/2794) of the down-regulated genes are supposed to be expressed at 8/16-cell stage (Fig 8b, 8c and S8b), indicating a ZGA defect. Strikingly, *KDM5s* were repressed in FK228-treated embryos (Fig 8d), and also resulted in an increase of H3K4me3 during ZGA (Fig 8e and 8f). In summary, these results suggested that HDAC1 has a conserved function in regulation of gene expression pattern during ZGA.

**Fig. 8.**
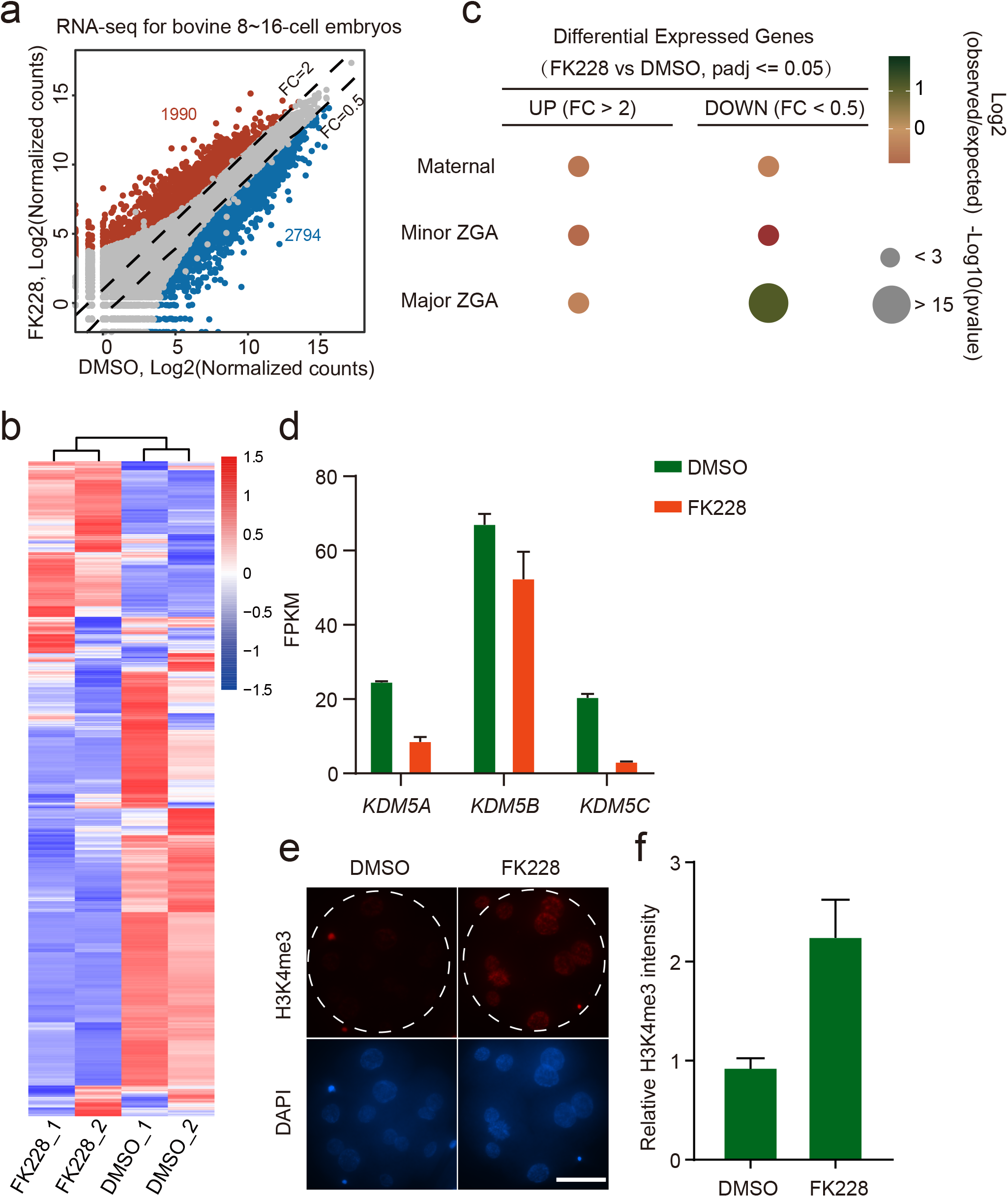
Function of HDAC1/2 on major ZGA is conservative in mouse and bovine embryos. **a,** Scatter plots showing gene expression level in embryos treated with DMSO (negative control) and FK228 in bovine 8/16-cell embryos. Two RNA-seq replicates are generated for differential expression analysis, and the read counts are normalized by DESeq2. Dash lines indicate the threshold of fold change (FK228/DMSO), and grey dots refer to genes with *padj* > 0.05, while dots in red and blue refer to genes with *padj* <= 0.05. Numbers of up- and down-regulated genes are also indicated in the figures. **b,** Relative expression of all bovine major ZGA genes in DMSO and FK228 treated embryos. **c,** Overlap of all differentially expressed genes with different gene categories. The gene categories are generated with *k-means* clustering of RNA-seq data for bovine MII oocytes and 4-cell, 8-cell, 16-cell embryos. The color of bubbles refers to log_2_ ratio of the number of observed genes in the gene set to randomly expected frequencies, and the size of bubbles shows -log_10_ of Fisher-exact test *p* value. **d,** Relative expression of *KDM5A, KDM5B, KDM5C* in DMSO and FK228 treated embryos. **e,f,** Immunofluorescence staining of H3K4me3 in bovine 8/16-cell embryos. e, representative images, scale bar, 50 μm, f, H3K4me3 intensity relative to DMSO-treated embryos.

**Fig. 9.**
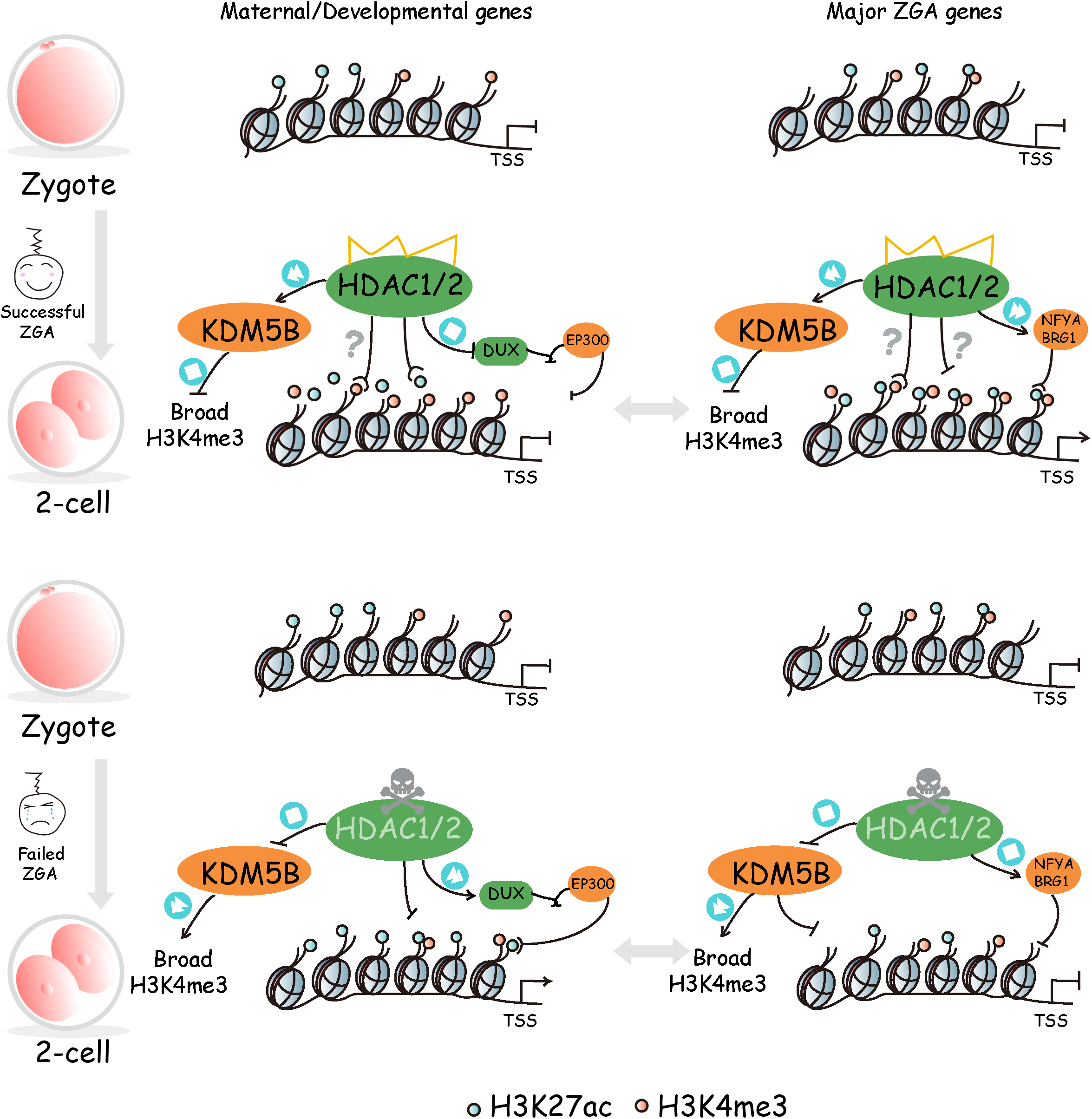
Model depicting the role of HDAC1/2 in ZGA. For maternal and developmental genes which are supposed to be silenced during ZGA, HDAC1/2 can deacetylated H3K27ac at both promoters and gene body regions. Concurrently, HDAC1/2 induces H3K4me3 accumulation at promoters indirectly and facilitates the expression of *Kdm5b*, which is responsible for the removal of broad H3K4me3 domain at distal regions. HDAC1 is also required for timely silencing of *Dux*, which otherwise induces expression of these silent genes. For actively expressed major ZGA genes, HDAC1/2 is required for H3K27ac and H3K4me3 enrichment at promoter regions, likely through an indirect way. Potentially, NFYA and BRG1 are also involved in HDAC1 effects on ZGA gene expression. However, HDAC1 still induce *Kdm5b* expression to remove broad H3K4me3 domain.

## DISCUSSION

A fundamental question in developmental biology is what factors trigger ZGA in mammals. Accumulative evidence revealed a growing list of proteins involved in the process^31, 32, 33, 34^. However, the molecular mechanisms of ZGA have yet to be determined. Here, we demonstrated HDAC1/2-mediated removal of H3K27ac is critical for establishing correct gene expression profile during ZGA. Mechanistically, HDAC1 regulates gene expression likely through controlling the locus-specific distribution of H3K27ac and H3K4me3. Importantly, HDAC1 acts as an upstream factor of KDM5B to regulate expression of target genes. We propose that HDAC1 is not only involved in development of the transcriptional repressive state for repressed genes but transcriptional active state for ZGA genes. To our knowledge, HDAC1/2 represent the first proteins hold a dual role in transcriptional regulation during ZGA. Histone acetylation has long been linked with transcriptional permissive state. In particular, H3K27ac is well established as a marker of active enhancers and promoters^21^. However, recent studies indicated H3K27ac is not necessary for transcriptional activity^39^. Whether H3K27ac results from transcriptional activation or it acts as a transcriptional inducer remains largely unknown especially during early embryogenesis, when dynamic chromatin reprogramming take place. Surprisingly, we observed a sharp decrease of global H3K27ac in both mouse and cattle. This phenomenon is not specific to H3K27ac and occurs to other histone acetylation during ZGA in mice, cattle, and pigs^24, 25^, suggesting histone deacetylation is a conserved event to set up chromatin state prior to ZGA. We speculated that H3K27ac reprogramming is one upstream inducer of the major ZGA since it occurs prior to the major ZGA and earlier than H3K4me3 reprogramming (early to late 2-cell stage), which appears as a ZGA regulator^8^.

The genome-wide histone deacetylation seems paradoxical when ZGA occurs. However, micro-scale ChIP-seq analysis indicate H3K27ac is not evenly decreased but undergo locus-specific reprogramming. In particular, H3K27ac is accumulated in the promoters of ZGA genes as expected. Moreover, it needs to be removed from the promoters of repressed genes, including developmental and maternal genes, suggesting H3K27ac deacetylation is required to develop the transcriptionally repressive state as described previously^1^. H3K27ac enrichment can be used to distinguish active from repressed genes compared with another two most well-addressed histone modifications (H3K4me3 and H3K27me3).

We demonstrated the histone deacetylase activity of HDAC1/2 is responsible for H3K27ac deacetylation and development progression throughout ZGA. In agreement with our data, previous reports show that HDAC1 is zygotically expressed beginning from 2-cell stage accompanying genome activation and only HDAC1 is sensitive to α-amanitin among the HDACs expressed in 2-cell embryos^15, 30^. HDAC2 seems more important during oogenesis while HDAC1 is an important transcriptional regulator in embryogenesis^15^. Upon fertilization, HDAC1 seems a key molecular player directly involved in development of transcriptional repressive state for maternal and developmental genes. Indeed, the dominant negative mutant of HDAC1 leads to up-regulated genes, the majority of them belong to maternal and developmental genes. Moreover, H3K27ac level are accumulated at promoter regions of up-regulated genes. Interestingly, we further identified DUX was enriched in H3K27ac-gained promoter regions. As an intron-less and multicopy gene that encodes a double-homeobox transcriptional factor involved in major ZGA, *Dux* is transiently transcribed during minor ZGA and its timely removal is critical for early embryogenesis^36^, of which the biological significance remains unclear. It is possible that the DUX removal is critical to inhibit the transcription activity at these repressed genes. Indeed, human DUX4 can recruit EP300 through its C-terminals and therefore involved in histone acetylation^37, 40^.

HDAC1/2 mutant elicit a dramatic downregulation of ZGA genes. These genes account for 24% of all ZGA genes. Grossly, above 50% ZGA genes are affected. Interestingly, H3K27ac was decreased at promoters of these genes after H1MU injection, in contrasting with HDAC1’s deacetylase activity. Thus, HDAC1 regulate transcriptional activity of these active genes in an indirect manner. It is possible that P300 is a core regulator of ZGA genes. P300 is under control of HDAC1 and we also documented a decrease of P300 in H1MU embryos (Table S2).

Crosstalk of different epigenetic modifications makes up precise regulatory network for gene expression^3, 41^. We observed that H3K27ac enrichment was much higher at promoters of H3K4me3 marked genes then those of H3K27me3 marked genes in mouse embryos, which conformed to the positive roles of H3K4me3 and H3K27ac in transcriptional activity. KDM5s were down-regulated in embryos injected with mutant *Hdac1* mRNA and caused inadequate removal of broad H3K4me3 in gene body and intergenic regions. Timely removal of broad H3K4me3 is viewed associated with mouse ZGA, but *Kdm5s*-knockdown embryos can develop beyond ZGA although a set of ZGA genes were slightly down-regulated at 2-cell stage^8, 9^. However, repression of HDAC1 could cause insufficient removal of H3K27ac and H3K4me3 concurrently, and led to embryos arrested at 2-cell stage. Besides, correcting H3K4me3 by overexpressing KDM5B could assist embryos overcoming ZGA block, suggesting that abnormal H3K4me3 alone plays limited roles in ZGA.

Altogether, we demonstrate an indispensable role for the histone deacetylase activity of HDAC1/2 during MZT in both mice and cattle. HDAC1 is required for the proper distribution of H3K27ac, and can also guide the removal of broad H3K4me3 indirectly. The chromatin reprograming is critical for the development of both transcriptional repressive state for silent genes and transcriptional active state for ZGA genes.

## MATERIALS AND METHODS

### Ethics Statement

All procedures involving laboratory animals were carried out in accordance with the guidelines for the care and use of laboratory animals and approved by Zhejiang University.

### Mouse embryo collection and in vitro culture

All experimental mice were raised and housed in a temperature controlled room (22-25°C) with a relative humidity of 60% in a 12-h light/dark cycle in Laboratory Animal Center at Zhejiang University. All the mice were provided access to food and water ad libitum. Eight to 10-week-old female BDF1 (C57BL/6 × DBA/2; Beijing Vital River Laboratory Animal Technology Co., Ltd.) mice were super-ovulated by an intraperitoneal injection of 7.5-10 IU PMSG (Sansheng Pharmaceutical Co. Ltd., Ningbo, China). Two days later, 7.5-10 IU hCG (Sansheng Pharmaceutical Co. Ltd., Ningbo, China) were administrated and mated with BDF1 male mice. 17-19 hrs after hCG injection, mice were sacrificed and zygotes were collected from the swollen upper part of the oviduct. Zygotes were incubated in the hyaluronidase (Sigma) solution to remove the cumulus cells and subsequently cultured in KSOM (Millipore) micro-drops under mineral oil at the condition of 37□ and 5% CO_2_.

### Bovine embryos *in vitro* production

Procedures of bovine *in vitro* production including *in vitro* maturation (IVM), *in vitro* fertilization (IVF) and *in vitro* culture (IVC) was carried out routinely in accordance with previously published studies^42, 43^. Briefly, cumulus-oocyte complexes (COCs) with >3 layers of cumulus cells were collected from ovaries derived from a local slaughterhouse. IVM was conducted with Medium-199 (M4530) supplemented with 10% FBS (Gibco-BRL, Grand Island, NY), 1 IU/ml FSH (Sansheng Biological Technology, Ningbo, China), 0.1 IU/ml LH (Solarbio, Beijing, China), 1 mM Na Pyruvate (Thermo Fisher Scientific, Waltham, MA), 2.5 mM GlutaMAX™ (Thermo Fisher Scientific, Waltham, MA), and 10 μg/mL Gentamicin. The IVM condition was 38.5°C under 5% CO_2_ in humidified air for 22-24 hrs. IVF was performed by incubating COCs (60-100 COCs per well in 4-well plates) with spermatozoa (1~5×10^6^), which was purified from frozen-thawed semen by using a Percoll gradient in BO-IVF medium (IVF bioscience, Falmouth, Cornwall, UK). IVF condition was 38.5°C under 5% CO_2_ for 9-12 hrs. Then, putative zygotes were removed of cumulus cells by pipetting up and down in Medium-199 (M7528) supplemented with 2% FBS (Gibco-BRL, Grand Island, NY). Embryos were cultured in BO-IVC medium (IVF bioscience, Falmouth, Cornwall, UK) at 38.5°C under 5% CO_2_ in humidified air until use.

### FK228 treatment experiments

Romidepsin (FK228, Depsipeptide, Selleck) dissolved in dimethyl sulfoxide (DMSO, Sigma) was added to KSOM (for mice embryo) or BO-IVC medium (for cattle embryo) at a final concentration of 50 nM. In control group, equivalent amount of DMSO (0.1%) was also added for embryo culturing.

### *In vitro* transcription and microinjection

Wild-type or mutant cDNA for *Hdac1, Hdac2* and *Kdm5b* were subcloned into T7-driven vectors. *Hdac1* mutant (H141A) and *Hdac2* mutant (H141A) were constructed as described previously ^29, 44^. All sequences were validated by Sanger sequencing prior to use (Sangon, Shanghai, China). To prepare mRNAs for microinjection, the plasmids were linearized and *in vitro* transcribed, capped and poly(A) tailed using T7 mMESSAGE mMACHINE Ultra Kit (Life Technologies, Grand Island, NY, USA) according to the manufacturer’s manual. RNA was recovered and purified by phenol:chloroform extraction and the integrity validated by gel electrophoresis.

mRNAs were microinjected into the cytoplasm of zygote using a Piezo-drill (Eppendorf, Germany) and Eppendorf transferman micromanipulators. *Hdac1* WT/mutant mRNA (500 ng/μl) or *Kdm5b* mRNA (1μg/μl) was loaded into microinjection needle and a constant flow was adjusted in order to achieve successful microinjection. Around 10 pl of mRNA was microinjected into the cytoplasm of zygotes or 2 cell blastomere.

### Immunofluorescence

Embryos were collected, washed and fixed for 10 minutes at room temperature in 4% formaldehyde in PBS (mouse: 10 min; cattle: 30 min). Embryo permeabilization was performed by treating with 0.5% Triton X-100 in PBS for 30 minutes in mice and 1 h in cattle. Then embryos were blocked in 10% fetal bovine serum in PBS for 1 h, and incubated in primary antibody for at least 1 h or 4 °C overnight. After 3 washes with 0.1% Triton X-100 in PBS, fixed embryos were incubated with secondary antibodies for at least 1 h. DNA was counterstained with DAPI to locate the nuclear region. Embryos were imaged with Zeiss LSM 880 confocal microscope. All the antibodies used in the present study were listed in Table S1.

### RNA-seq library preparation and sequencing

Mouse early 2-cell and late 2-cell stage embryos (50 embryos/sample, n=2 biological replicates/group) were collected at 24 h and 36 h post fertilization, respectively. Bovine 8/16-cell stage embryos (30 embryos/sample, n=2) were collected at 72 h post fertilization. Total RNA was extracted using Arcturus PicoPure RNA Isolation Kit (Life Technologies, Grand Island, NY, USA) according to the manufacturer’s manual. Then mRNAs were separated using oligo(dT)25 beads. Sequencing libraries were constructed by using NEB Next Ultra RNA Library Prep Kit for Illumina (E7530) based on the instruction. Briefly, mRNAs were fragmented and reverse transcribed. cDNA library underwent end repair, poly(A)-tailing, adaptor ligation, and PCR amplification for 12–15 cycles in order to prepare the sequencing libraries. Paired-end 150 bp sequencing was performed on a NovaSeq (Illumina) platform by Novogene Co., Ltd.

### ULI-NChIP-seq library preparation and sequencing

Mouse late 2-cell embryos (120 embryos/sample, n=2) were collected at 48 hrs after hCG injection. The zona pellucidae of the embryos were removed with Acid Tyrode’s solution, then the embryos were washed 3 times in 0.5% bovine serum albumin (Millpore) in DPBS (Gibco) before flash-freezing in liquid nitrogen. ULI-NChIP was performed according to the published protocols with slight modifications^12^. One microgram of H3K4me3 (Cell Signaling Technology, #9751) or H3K27ac (Active Motif, AM39133) antibody was used for each immunoprecipitation reaction. ULI-NChIP-seq libraries were generated using the NEB Ultra DNA Library Prep Kit (E7645) according to the manual. Paired-end 150 bp sequencing was performed on a NovaSeq (Illumina) platform by Novogene Co., Ltd.

### RNA-seq data alignment and quantification

The raw sequencing reads were trimmed with Trimmomatic (version 0.39)^45^ to get rid of low-quality reads and adaptor sequences. Clean reads were then mapped to mm10 (mouse) or ARS-UCD1.2 (bovine) with Hisat2 (version 2.1.0)^46^. The raw read counts of genes were calculated with featureCounts (version 1.6.3)^47^. For quantification of repetitive element, repeat annotations were downloaded from UCSC genome browser, and the raw counts of repetitive element were counted with BEDTools (version 2.30.1) ^48^. Gene expression values were normalized to FPKM with Cufflinks (version 2.2.1) ^49^ for heatmap, line plot and bar plot visualization. The raw counts of repetitive elements were normalized with DESeq2^50^ and then used for box pot visualization.

### Differential expression analysis and functional annotation

Reads counts from FeatureCounts (genes) or BEDTools (repetitive elements) were used for differential expression analysis with DESeq2. The differentially expressed genes or repetitive elements between groups were identified when fold change >2 or <0.5 and adjusted *P* value < 0.05. The differential expression of repetitive elements was further validated by TEtranscripts python package (version 2.2.1)^51^. Functional annotation and enrichment analysis of differentially expressed genes was performed with the Database for Annotation, Visualization and Integrated Discovery (DAVID)^52, 53^.

### Classification of gene sets and clustering analysis

RNA-seq data of mouse and bovine oocytes and embryos was obtained from Jingyi Wu *et al.* ^26^and Alexander Graf *et al*^54^, respectively. The FPKM matrixes were generated with Cufflinks as described in RNA-seq data alignment and quantification section. Then the non-zero FPKM of genes was used for *k*-means clustering, which was performed with R based on Euclidean distance. After *k*-means clustering, genes with similar expression pattern were classified as maternal, minor ZGA, major ZGA or developmental genes. Clustering of differentially expressed genes in Fig. 3d was also performed as above. For hierarchical clustering of mouse late 2-cell embryos in different groups (Fig. S4e), euclidean distance of samples was calculated based on FPKM matrixes and hierarchical clustering was performed with hclust (dist, method = ‘average’) in R (http://www.rproject.org).

### ChIP-seq data processing

The raw sequencing reads were trimmed with Trimmomatic (version 0.39)^45^ to remove residual adapter sequences and low-quality reads. Then the clean reads were aligned to mm10 using Bowtie2 (version 2.3.5)^55^ with default parameters. Alignments with low mapping quality were discarded by SAMtools (version 1.7)^56^, and PCR duplicates were removed with Picard (version 2.23; https://broadinstitute.github.io/picard/). To visualize H3K4me3 and H3K27ac signal in IGV genome browser, the genome was binned into 50 bp windows and RPKM for each window was calculated using bamCoverage function from DeepTools^57^.

### Correlation between biological replicates

The genome was binned into 2000 bp windows using makewindows utility of BEDTools^58^, and the reads counts of each window were calculated and normalized to RPKM. Then the RPKM value for each sample was used to calculate Pearson correlation coefficient and draw scatter plots with R (http://www.rproject.org).

### ChIP-seq peak calling

The peak number is affected by sequencing depth and equal numbers of ChIP and input reads result in best performance of peak callers^59, 60^. Therefore, we merged alignments of biological replicates and randomly subsampled them to equivalent depth using SAMtools before peak calling. The peaks were called by MACS2 (version 2.2.7.1)^61^ using the following parameters: -B -p 1e-5 --nomodel --broad --extsize 73. The peaks with fold enrichment less than 2 were removed. The filtered peaks were then annotated using ChIPseeker^62^.

### Identification of H3K27ac and H3K4me3-gained or -lost regions

To compared H3K27ac or H3K4me3 signals between groups, the mouse genome was scanned using a sliding window of 5 kb and step size of 1 kb, and then RPKM for each window was calculated. Next, H3K27ac or H3K4me3 signals were compared parallelly between H_2_O and *Hdac1* MU. The H3K27ac and H3K4me3-gained or -lost regions were identified with following threshold: log_2_ (fold change) > 1.5 for gained regions or < −1.5 for lost regions, and sum of RPKM in H_2_O and *Hdac1* MU >1. Then the selected regions were merged with BEDTools if they were overlapped, and the genome coverage of merged regions was calculated by genomecov mode of BEDTools.

### Motif analysis

The mouse genome was first scanned using a sliding window of 1 kb and step size of 1 kb, and then RPKM of H3K27ac for each window was calculated. Next, the ratio of H3K27ac signal in *Hdac1* MU to H_2_O was calculated, and windows with top 2% of ratios were retained as the most increased H3K27ac regions. On the other hand, windows with top 2% of ratios of H_2_O to *Hdac1* MU were retained as the most decreased H3K27ac regions. The selected increased or decreased regions were merged with BEDTools if they were overlapped. The merged regions were used for motif identification by Homer (version 4.11; http://homer.ucsd.edu/homer/motif/) with the parameters: -size given -p 10 -len 8.

### Datasets

Previously published datasets used in the present work include H3K27ac ChIP-seq (GSE72784), H3K4me3 ChIP-seq (GSE71434), Mouse RNA-seq (GSE66582), and Bovine RNA-seq (GSE52415). Our RNA-seq and ULI-NChIP-seq data have been deposited to GEO with the accession number GSE182555.

## Acknowledgments

We thank Dr. Li Shen, Dr. Xudong Fu and Dr. Wei Xie for their helpful discussions. Thank Dr. Xukun Lu from Dr. Wei Xie’s lab and Dr. Zhe Zhang from Yuchun Pan’s lab for their help on bioinformatic analysis. This project was supported by National Natural Science Foundation of China (No. 31672416, No. 31872348, and No. 32072731 to K.Z.; No.31941007 to L.L. and S.W.; No. 32072939 to H.W.) and Zhejiang Provincial Natural Science Foundation (LZ21C170001 to K.Z. and No. LY19C180002 to H.W.) and China Postdoctoral Science Foundation (No. 2020M671742 to L.L.).

